# Metagenomic- and cultivation-based exploration of anaerobic chloroform biotransformation in hypersaline sediments as natural source of chloromethanes

**DOI:** 10.1101/858480

**Authors:** Peng Peng, Yue Lu, Tom N.P. Bosma, Ivonne Nijenhuis, Bart Nijsse, Sudarshan A. Shetty, Alexander Ruecker, Aleksandr Umanetc, Javier Ramiro-Garcia, Andreas Kappler, Detmer Sipkema, Hauke Smidt, Siavash Atashgahi

## Abstract

Chloroform (CF) is an environmental contaminant that can be naturally formed in various environments ranging from forest soils to salt lakes. Here we investigated CF removal potential in sediments obtained from hypersaline lakes in Western Australia. Reductive dechlorination of CF to dichloromethane (DCM) was observed in enrichment cultures derived from sediments of Lake Strawbridge, which has been reported as a natural source of CF. The lack of CF removal in the abiotic control cultures without artificial electron donors indicated that the observed CF removal is a biotic process. Metabolite analysis with ^13^C labelled CF in the sediment-free enrichment cultures (pH 8.5, salinity 5%) revealed that increasing the vitamin B_12_ concentration from 0.04 to 4 μM enhanced CF removal, reduced DCM formation, and increased ^13^CO_2_ production, which is likely a product of CF oxidation. Known organohalide-respiring bacteria and reductive dehalogenase genes were neither detected by quantitative PCR nor metagenomic analysis. Rather, members of the order *Clostridiales*, known to co-metabolically transform CF to DCM and CO_2_, were detected in the enrichment cultures. Genome-resolved metagenome analysis indicated that their genomes encode enzymatic repertoires for the Wood-Ljungdahl pathway and cobalamin biosynthesis that are known to be involved in co-metabolic CF transformation.

**Importance:** More than 90% of the global CF emission to the atmosphere originates from natural sources, including saline environments such as salt lake sediments. However, knowledge about the microbial metabolism of CF in such extreme environments is lacking. Here we showed CF transformation potential in a hypersaline lake that was reported as a natural source of CF production. Application of interdisciplinary approaches of microbial cultivation, stable isotope labelling, and metagenomics aided in defining potential chloroform transformation pathways. This study indicates that microbiota may act as a filter to reduce CF emission from hypersaline lakes to the atmosphere, and expands our knowledge of halogen cycling in extreme hypersaline environments.

## Introduction

Until the 1970s, halogenated organic compounds, organohalogens, were believed to originate exclusively from anthropogenic sources (1). This long-held view was changed following the discovery of diverse organohalogens from natural environments. To date, over 5000 naturally occurring organohalogens have been identified (2). A remarkable example is chloroform (trichloromethane, CF) which is a known environmental contaminant and a potential carcinogen that bioaccumulates and is harmful for living organisms (3). CF is synthetically produced in chemical industries as an anesthetic, as an intermediate for the production of refrigerants, and as a degreasing agent and fumigant (4). However, anthropogenic sources were estimated to contribute to less than 10% of the annual 700,000–820,000 tons global CF production (5). Natural CF emissions have been reported from numerous terrestrial and aquatic environments such as forest soils (6–9), rice fields (10), groundwater (11), oceans (12), and hypersaline lakes (13, 14). Biotic and abiotic processes like burning of vegetation, chemical production by reactive Fe species, and enzymatic halogenation can lead to natural production of CF (15). Similar to other low molecular weight volatile organohalogens (VOX), CF release into the atmosphere can cause ozone depletion and impact climate change (16).

CF is persistent in the environment and is hardly dechlorinated/degraded under oxic conditions due to the three chlorine substitutes (17, 18). In contrast, microbial CF transformation is often mediated by anaerobic microbes (19–23). Anaerobic CF transformation has been reported to be mediated by acetogens like *Acetobacterium woodii* (24) and *Clostridium* sp. (25), methanogenic *Methanosarcina* spp. (26–28), and fermentative *Pantoea* spp. (23) producing dichloromethane (DCM), carbon monoxide (CO) and/or carbon dioxide (CO_2_). This is a co-metabolic process for which the responsible genes and enzymes are not yet clear. Previous studies indicated that co-metabolic CF transformation was likely mediated by enzymes involved in the Wood-Ljungdahl pathway and methanogenesis (24, 29). Moreover, transition-metal co-factors, e.g. cob(I)/cob(II)alamins and F_430_ (nickel(I)-porphinoid), that facilitate key enzymes of acetogenesis (5-methyltetrahydrofolate corrinoid/iron-sulfur protein methyltransferase) and methanogenesis (methyl-coenzyme M reductase) can act as reductants and nucleophilic reagents catalyzing nonspecific reductive dechlorination of chloromethanes (30–32).

Another group of anaerobes known as organohalide-respiring bacteria (OHRB) can use CF as a terminal electron acceptor, and couple CF reductive dechlorination to energy conservation (33, 34). For instance, CF respiration to DCM has been reported using *Desulfitobacterium* sp. strain PR (35), *Desulfitobacterium hafniense* TCE1 (36), *Dehalobacter* sp. strain UNSWDHB (37, 38) and a mixed culture containing *Dehalobacter* (21). The enzymes responsible for reductive dehalogenation in OHRB are mainly corrinoid cofactor dependent reductive dehalogenases (RDases) such as a CF RDase (CfrA) identified from *Dehalobacter*-containing microbial consortia (39). CF can also be abiotically dechlorinated under anoxic conditions via hydrogenolysis to DCM, or via reductive elimination to CH_4_ (40–42).

Previous studies have shown the presence of organohalogen-metabolizing microbes in environments where natural organohalogens have been shown or suspected to be present (43, 44). Hypersaline lakes are natural sources of VOX, and (micro)organisms are major contributors of VOX emission in these environments (13, 14, 45). Moreover, NaCl in hypersaline lakes might promote high rates of organic matter halogenation (46). However, knowledge about the microbial metabolism of VOX in such extreme environments is lacking. This information is necessary to understand whether microbes can act as a filter for VOX in hypersaline environments that at least partly prevent their release to the atmosphere. In this study, we prepared microcosms from the sediments of hypersaline Lake Strawbridge and Lake Whurr in Western Australia. Lake Strawbridge has been reported as a natural source of chloromethane (CM) and CF (13). The CF transformation process and responsible microbes were studied by a combination of anaerobic cultivation, stable isotope labelling, molecular analyses, and metagenomics.

## Results

### Physical-chemical characteristics of sediments

The top (0-12 cm) and bottom (>12 cm) layer sediments of Lake Strawbridge were slightly alkaline with a pH ranging from 8.2 to 8.5 whereas those of Lake Whurr were acidic with a pH of 4.6–5.4 (Table 1). Salinity, water content and total organic carbon (TOC) were higher in the top layer compared to the bottom layer for both lake sediments (Table 1). Sodium (17.5–71.1 mg/g dry sediment) and chloride (31.9–123.5 mg/g dry sediment) were dominant among the cations and anions, respectively. Nitrate and chlorate were neither detected in top- nor in bottom-layer sediments (Table 1).

**Table 1.** Geochemical properties of Lake Strawbridge and Lake Whurr sediments. Duplicates sediment cores from each hypersaline lake are labelled as as LS1&LS2 and LW1&LW2. TOP :0-12 cm depth, BOT: >12 cm depth

|  | Lake Strawbridge (LS) |  |  |  | Lake Whurr (LW) |  |  |  |
| --- | --- | --- | --- | --- | --- | --- | --- | --- |
|  | LS1-TOP | LS2-TOP | LS1-BOT | LS2-BOT | LW1-TOP | LW2-TOP | LW1-BOT | LW2-BOT |
| pH <sup>a</sup> | 8.2 | 8.3 | 8.5 | 8.5 | 5.4 | 5.4 | 4.5 | 4.6 |
| Water content (%) | 37.3 | 27.3 | 16.7 | 15.4 | 26.0 | 25.7 | 24.2 | 23.0 |
| Salinity (%) | 17 | 14 | 5 | 5 | 15 | 20 | 11 | 11 |
| TOC (g/kg dry sediment) | 21 | 15 | 5 | 5 | 12 | 14 | 6 | 6 |
| Na (mg/g dry sediment) | 57.0 | 48.5 | 17.5 | 18.1 | 55.0 | 71.1 | 35.0 | 35.8 |
| Ca (mg/g dry sediment) | 0.7 | 0.8 | 0.1 | 0.2 | 6.8 | 4.2 | 0.3 | 0.3 |
| K (mg/g dry sediment) | 2.0 | 2.0 | 1.0 | 0.9 | 1.7 | 1.8 | 1.1 | 1.2 |
| Mg (mg/g dry sediment) | 2.8 | 2.9 | 1.1 | 1.1 | 4.5 | 4.6 | 3.5 | 3.4 |
| Total Fe (mg/g dry sediment) | 6.5 | 6.3 | 2.2 | 1.9 | 1.5 | 3.2 | 0.3 | 0.6 |
| Cl<sup>-</sup> (mg/g dry sediment) | 101.3 | 84.7 | 31.9 | 33.1 | 93.1 | 123.5 | 64.8 | 64.0 |
| SO<sub>4</sub><sup>2-</sup> (mg/g dry sediment) | 3.9 | 3.6 | 1.5 | 1.8 | 19.6 | 14.8 | 4.3 | 4.4 |
| NO<sub>3</sub><sup>-</sup> (mg/g dry sediment) | n.d. | n.d. | n.d. | n.d. | n.d. | n.d. | n.d. | n.d. |
| ClO<sub>3</sub><sup>-</sup> (mg/g dry sediment) | n.d. | n.d. | n.d. | n.d. | n.d. | n.d. | n.d. | n.d. |

<sup>a</sup> Measured in 0.01 M CaCl<sub>2</sub> after 2 h
n.d. not detected

### CF dechlorination in enrichment cultures

No CF dechlorination was observed in the enrichment cultures with the sediment from Lake Whurr after 70 days of incubation (data not shown), whereas CF was reductively dechlorinated to DCM in the enrichment cultures with the sediment from Lake Strawbridge (Fig. 1A–D). CM and methane as potential products of CF transformation were not detected (data not shown), despite an evident lack in the mass balance between CF disappearance and DCM production in sediment cultures and some transfer cultures (Fig. 1A–F). The lack of methane production also suggested inhibition and/or absence of methanogens. The fastest CF dechlorination rate (1.82 μmol/day/L) to DCM was observed in the enrichment cultures with the bottom layer sediments from Lake Strawbridge in the modified growth medium (MGM) medium (Fig. 1B). Therefore, these cultures were selected to obtain sediment-free cultures in subsequent transfers (Fig. 1E–G).

**Fig. 1.**
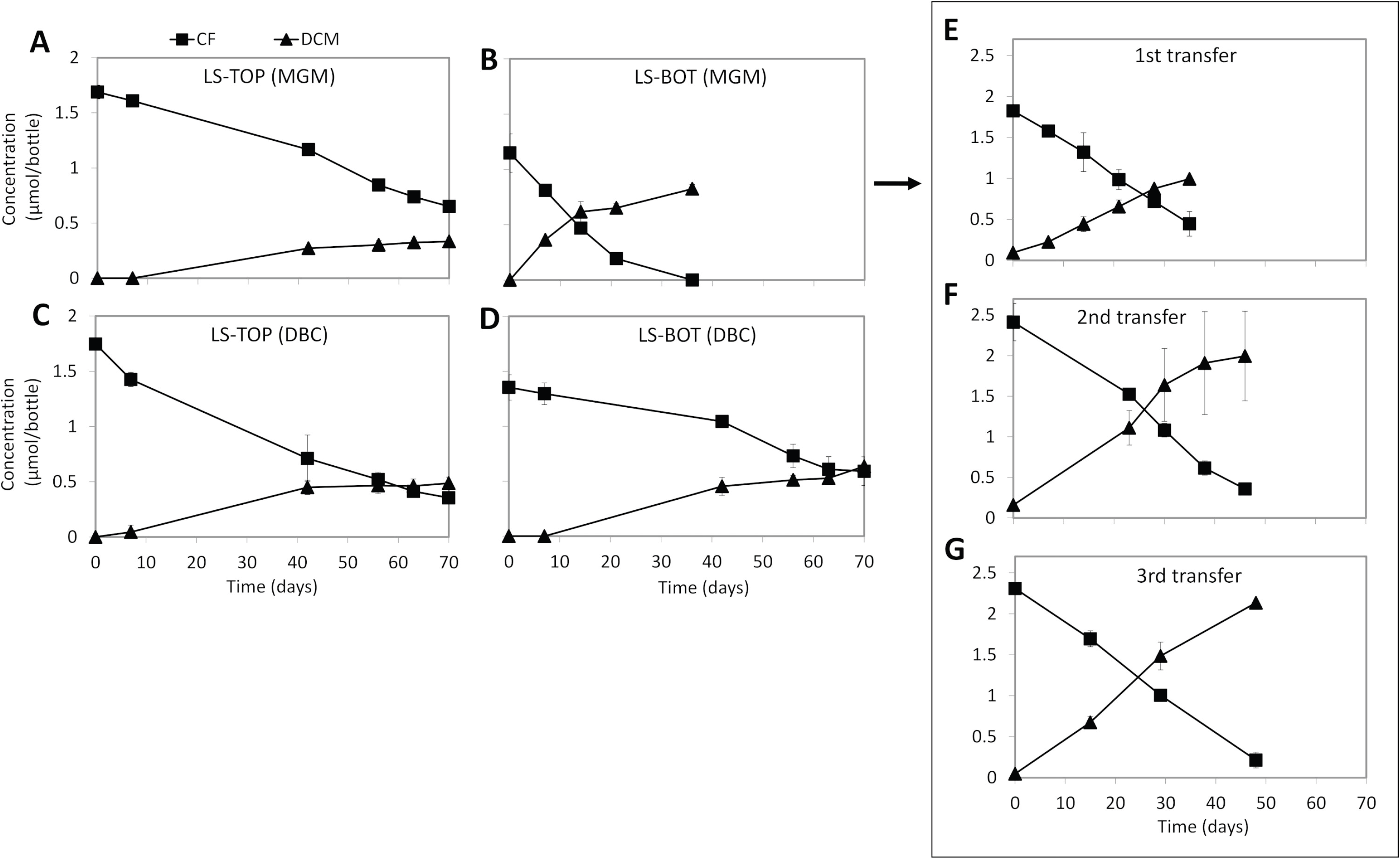
CF transformation in the sediment enrichment cultures and subsequent transfer cultures. Dechlorination of CF in MGM with top layer (LS-TOP, A) and bottom layer sediments (LS-BOT, B) from Lake Strawbridge, and dechlorination of CF in DBC medium with top (C) and bottom layer (D) sediments from the same lake. Dechlorination of CF in subsequent transfer cultures of the bottom layer sediment enrichment cultures with MGM (E, F, G). Points and error bars represent the average and standard deviation of samples taken from duplicate cultures.

Adding vitamin B_12_ from 0.04 to 4 μM steadily increased CF dechlorination rates in the sediment-free cultures (Fig. 2). For instance, in the cultures amended with 4 μM vitamin B_12_, the CF dechlorination rate reached 31.9 μmol/day/L (Fig. 2E), ~30 times higher than the dechlorination rate in the cultures without extra vitamin B_12_ supplementation (~0.9 μmol/day/L) (Fig. 1E–G). In turn, increasing vitamin B_12_ concentration led to concurrent decrease of DCM accumulation. Accordingly, less than 30% of the CF was converted to DCM in the cultures amended with 4 μM vitamin B_12_ (Fig. 2E). No CF dechlorination was observed in the abiotic controls even in the presence of 4 μM vitamin B_12_ (data not shown). In contrast, CF dechlorination to DCM and (or) CM was observed in abiotic controls when either Ti(III) or dithiothreitol (DTT) were used as an artificial electron donor together with 4 μM vitamin B_12_ (Fig. S1).

**Fig. 2.**
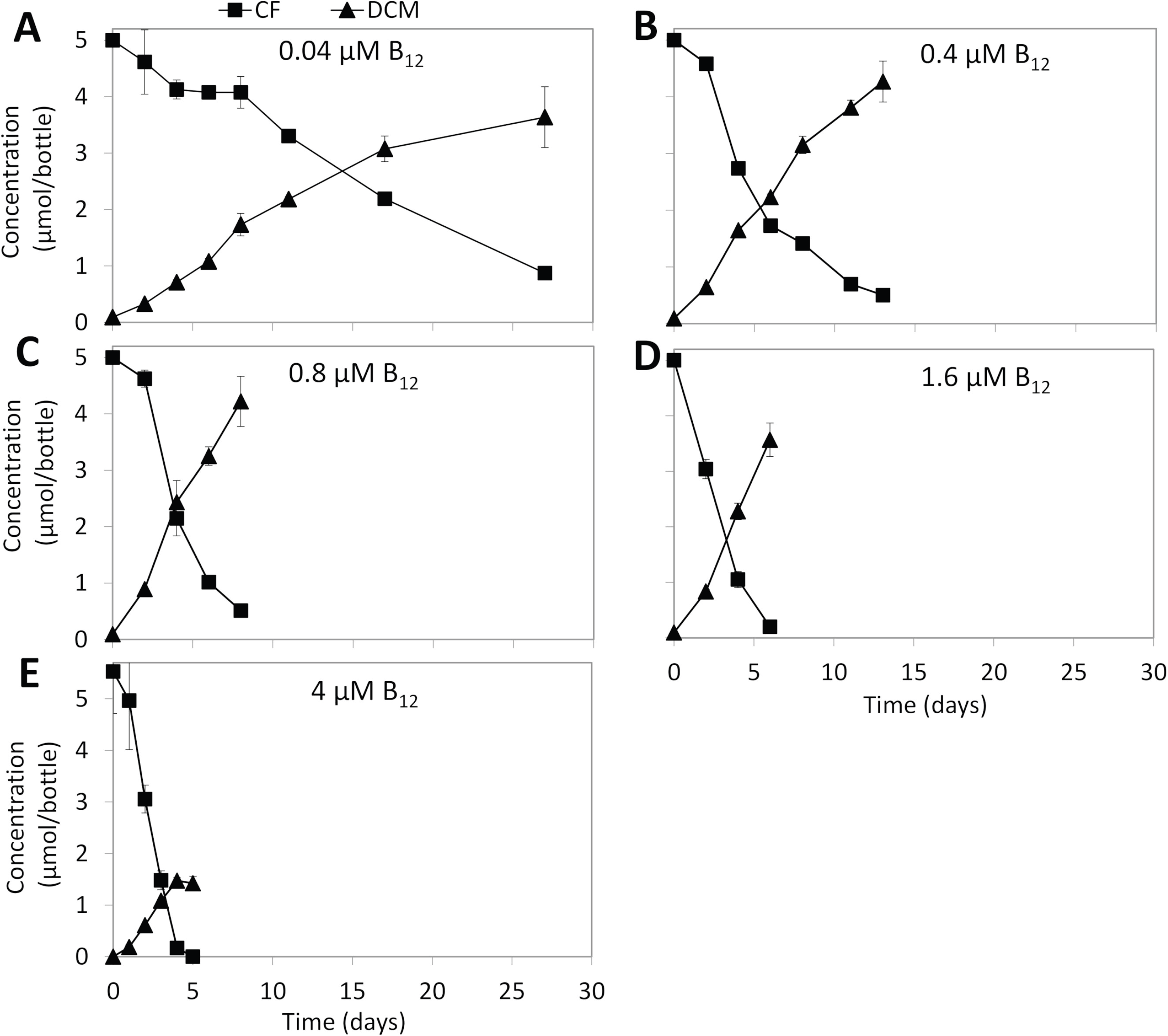
CF transformation in sediment-free cultures amended with 0.04 (A), 0.4 (B), 0.8 (C), 1.6 (D), and 4 μM (E) vitamin B_12_. Points and error bars represent the average and standard deviation of samples taken from duplicate cultures.

### Analysis of ^13^CO_2_ production from ^13^CF

^13^CO_2_ production was detected in the culture containing 1.25 μmol/bottle ^13^C-labelled CF, 3.75 μmol/bottle non-labelled CF and 4 μM vitamin B_12_ during the incubation (Fig. 3A). Recovery of ^13^C to ^13^CO_2_ was only detected in the culture with ^13^C-labelled CF as indicated by the increasing of *δ*^13^C value from −23.42‰ to 263.46‰ during the incubation (Fig. 3B). At day 5, 0.84 μmol/bottle ^13^CO_2_ and 1.7 μmol/bottle DCM were detected (Fig. 3A). Assuming that 25% of the DCM (0.43 μmol/bottle) originated from ^13^C-labelled CF (comprising 25% of total CF mass), a ca. 100% ^13^C conversion of CF to CO_2_ and DCM as the main products can be envisaged where removal of 1.25 μmol/bottle ^13^C-labelled CF resulted in production of 0.43 μmol/bottle ^13^C-DCM and 0.84 μmol/bottle ^13^CO_2_.

**Fig. 3.**
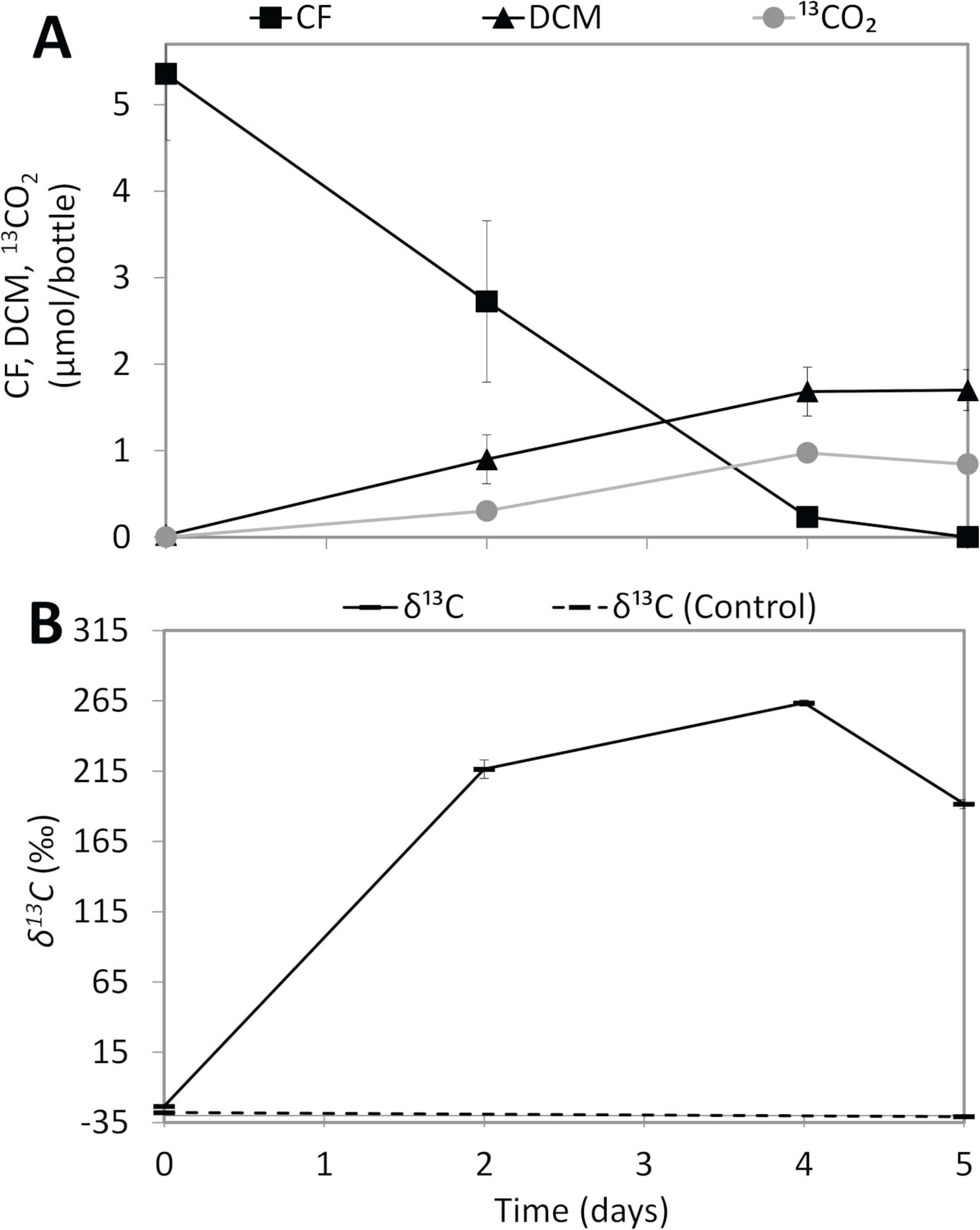
^13^CO_2_ production from CF (A) and δ^13^C values (B) in the sediment-free cultures amended with 1.25 μmol/bottle ^13^C-labelled CF, 3.75 μmol/bottle non-labelled CF, and 4 μM vitamin B_12_. Control cultures contained the same concentrations of non-labelled CF and vitamin B_12_. Points and error bars represent the average and standard deviation of samples taken from duplicate cultures.

### qPCR and bacterial community analysis

Bacterial and archaeal 16S rRNA gene copies in the top sediment layers of Lake Whurr and Lake Strawbridge were at least one order of magnitude higher than the 16S rRNA gene copies in bottom layers of the same lakes (Fig. S2A). The top layer sediment from Lake Strawbridge had the highest number of 16S rRNA gene copies of bacteria [(3.3 ± 0.87) × 10^8^ copies/g dry sediment] and archaea [(8.6 ± 0.25) × 10^7^ copies/g dry sediment] among all the sediments from the two lakes (Fig. S2A). Sediment enrichment cultures and subsequent transfer cultures prepared from the bottom layer sediment of Lake Strawbridge, contained 10^6^–10^7^ bacterial 16S rRNA gene copies/ml culture (Fig. S2B). However, archaeal 16S rRNA gene copies decreased dramatically to ~10^4^ copies/ml in the sediment enrichment cultures, and to below 10^2^ copies/ml culture in the transfer cultures (Fig. S2B). Known OHRB including *Desulfitobacterium*, *Dehalobacter*, *Dehalococcoides*, *Geobacter* and *Sulfurospirillum* were not detected in the enrichment cultures (data not shown).

Bacterial community analysis based on Illumina sequencing of barcoded 16S rRNA gene V1–V2 region amplicons showed that *Cyanobacteria*, *Chloroflexi*, *Proteobacteria* and *Firmicutes* were the most abundant phyla (cumulative relative abundance > 70%) in top and bottom layer sediments of Lake Strawbridge (Fig. S3). The relative abundance of *Clostridiales* and *Halanaerobium* (*Firmicutes*) increased from 5–16% (*Clostridiales*) and 3–7% (*Halanaerobium*) in the bottom layer sediments to ~67% and ~18%, respectively, in the initial and subsequent transfer enrichment cultures (Fig. S3). The relative abundance of *Desulfovibrio*, a genus within the *Proteobacteria*, increased from less than 0.1% to 0.3–8% in the initial and subsequent transfer enrichment cultures.

### Metagenomic analysis

Binning of the assembled metagenome sequences allowed the reconstruction of six near complete (>95%) metagenome-assembled genomes (MAG) of *Clostridiales* (bin1, bin3, bin4), *Halanaerobium* (bin2), *Bacillus* (bin5) and *Desulfovibrio* (bin6) (Table S1) accounting for ~84–95% relative abundance in sediment-free cultures with and without B_12_ (Fig. 4A). *Desulfovibrio* (bin6) was the most abundant MAG (45% relative abundance) in the cultures without external B_12_ addition, which is different than the results obtained from 16S rRNA gene based bacterial community analysis (0.4–9% relative abundance) (Fig. S3). One reason for this might be the change of the growth medium components in the sediment-free cultures used for metagenomic analysis that contained glycerol as carbon source and lower amounts of yeast extract and peptone compared to the cultures used for 16S rRNA gene analysis (Table S2, Fig. 4A). Except for bin 4, the addition of vitamin B_12_ (4 μM) increased the relative abundance of *Firmicutes* MAGs.

**Fig. 4.**
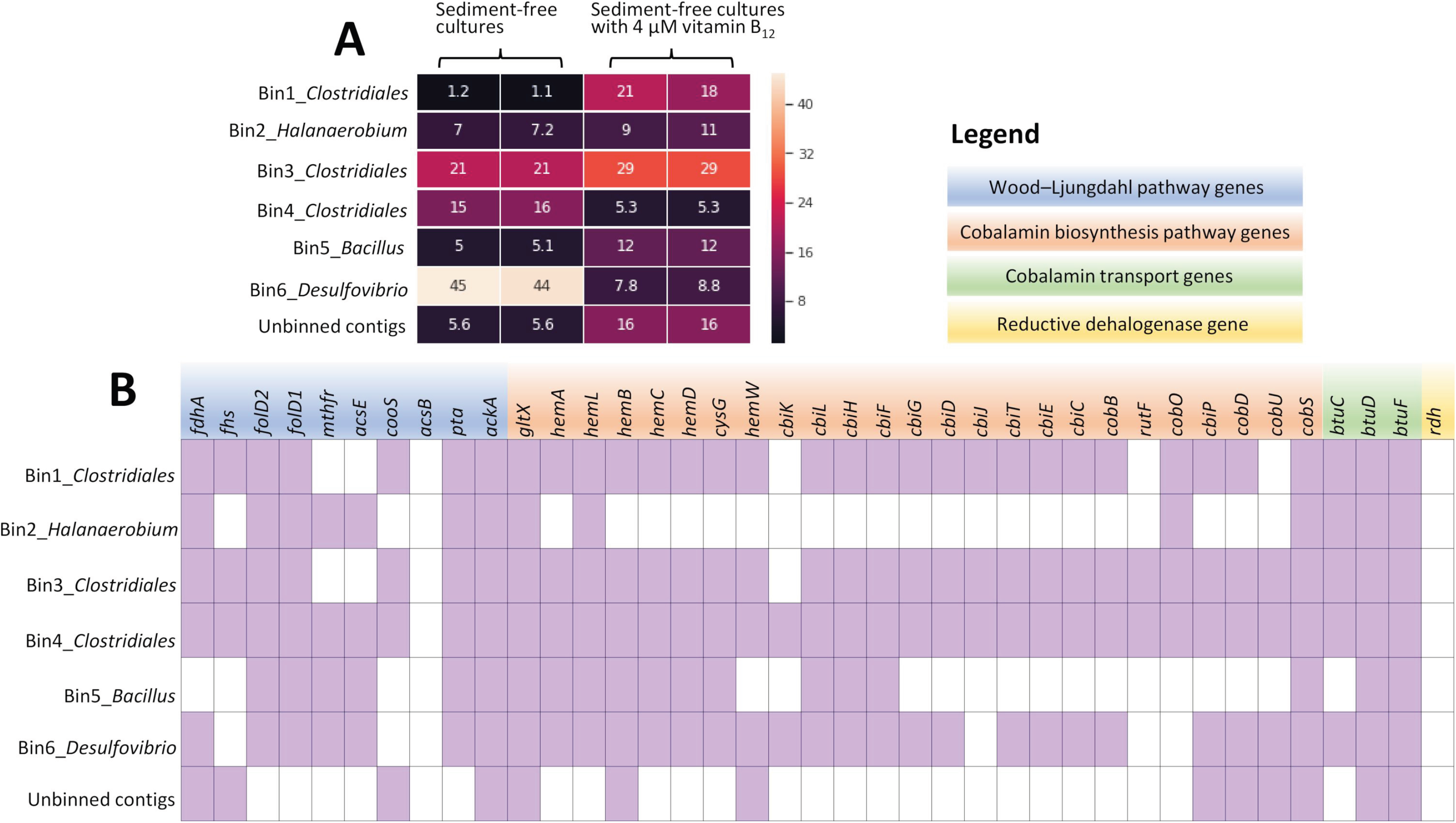
Heatmap of relative abundance of the MAGs and unbinned contigs assembled from metagenomes of the sediment-free enrichment cultures with and without addition of 4 μM vitamin B_12_ (A), and presence and absence of genes involved in the Wood–Ljungdahl pathway, cobalamin biosynthesis and transport and reductive dehalogenation (organohalide respiration) in the MAGs and unbinned contigs (B).

Reductive dehalogenase genes (*rdh*) were neither detected in the MAGs nor the unbinned contigs (Fig. 4B). In contrast, most of the genes encoding enzymes from the Wood-Ljungdahl pathway and cobalamin biosynthesis, all of which have been suggested to be involved in co-metabolic chloroform transformation, were identified in different MAGs (Fig. 4B). A notable exception was the absence of the *acsB* gene encoding acetyl-CoA synthase, the signature gene of the Wood-Ljungdahl pathway (Fig. 4B).

## Discussion

The present study showed CF dechlorination to DCM and CO_2_ in microcosms with sediments from hypersaline Lake Strawbridge in Western Australia, which has previously been shown to be a natural source of CF and CM (13). The lack of CF removal in the abiotic control cultures without artificial electron donors (Ti(III) or DTT) indicated that the CF removal in the sediment and sediment-free enrichment cultures is a biotic process and at least needs cellular metabolism for electron donor generation. Known CF-respiring bacteria such as *Desulfitobacterium* (35, 36) and *Dehalobacter* (38) were neither detected in the sediment microcosms nor in the sediment-free cultures by qPCR (data not shown) or 16S rRNA gene-targeted bacterial community analysis (Fig. S2, S3). Moreover, *rdh* genes were not detected in any of the MAGs or unbinned contigs, indicating that CF respiration by OHRB was unlikely.

Compared to the original sediment samples used for preparation of the microcosms, relative abundance of the order *Clostridiales* was significantly increased in the sediment and sediment-free enrichment cultures (Fig. S3). Acetogens belonging to this order such as members of the genera *Clostridium* and *Acetobacterium* have previously been shown to mediate co-metabolic CF dechlorination (24, 25, 47). For instance, *Clostridium* sp. strain TCAIIB was shown to dechlorinate CF to DCM and unidentified products (25), although underlying mechanisms remain unknown. One plausible explanation can be conversion of vitamin B_12_ (cob(III)alamin) to cob(I)/cob(II)alamins by *Clostridium* species (48, 49) that can mediate CF dechlorination.

Addition of extra vitamin B_12_ shifted the dominant CF transformation pathway from reductive dechlorination to DCM, to oxidation to CO_2_ (Fig. 2, 3). This is in line with previous studies using fermentative (23) and methanogenic enrichment cultures (20, 22, 50). CF oxidation was proposed to occur via the net hydrolysis of CF to CO (23, 24), but the enzymes responsible for such a transformation have not been identified. Another study suggested a possible role of vitamin B_12_-dependent Wood-Ljungdahl pathway enzyme(s) in CF hydrolysis to CO (24). However, CF hydrolysis to CO was also observed in non-acetogenic and fermentative *Pantoea* spp. amended with vitamin B_12_ (23), indicating a possible role of other (vitamin B_12_-dependent) pathways in CF hydrolysis. Except for the acetyl-CoA synthase gene, we detected all genes encoding enzymes involved in the Wood-Ljungdahl pathway as well as genes for cobalamin (enzymatic cofactor for methylenetetrahydrofolate reductase (MTHFR)) biosynthesis and transport in the *Clostridiales* MAGs (Fig. 4B). However, considering the slower CF transformation in the sediment-free cultures (Fig. 1F) as opposed to the original cultures (Fig. 1B), these MAGs were not likely (the main) vitamin B_12_ producers. Vitamin B_12_ producing microorganisms likely decreased in the sediment-free enrichment cultures during the enrichment process (Fig. S2, S3). In turn, the amount of CF (2―5 μmol/bottle or 50―100 μM) added in the enrichment cultures might have exceeded the toxic level for many vitamin B_12_ producing bacteria and archaea. CF is known highly toxic for some bacteria and archaea, and growth inhibition of acetogenic bacteria and methanogenic archaea was noted at concentrations as low as 0.1 μM (20). Previous studies using the sediments of lake Strawbridge reported natural CF production of ~0.017 μmol/kg dry sediment (13), that may exert a negligible inhibitory effect on the vitamin B_12_ producing microorganisms, suggesting a high molar ratio of vitamin B_12_ to CF in the sediment that may mediate CF conversion to CO/CO_2_. The CO produced from CF could be further oxidized to CO_2_ by CO dehydrogenase (CooS, Fig. 5) (50). We did not detect CO in the enrichment cultures likely due to its rapid conversion to CO_2_.

**Fig. 5.**
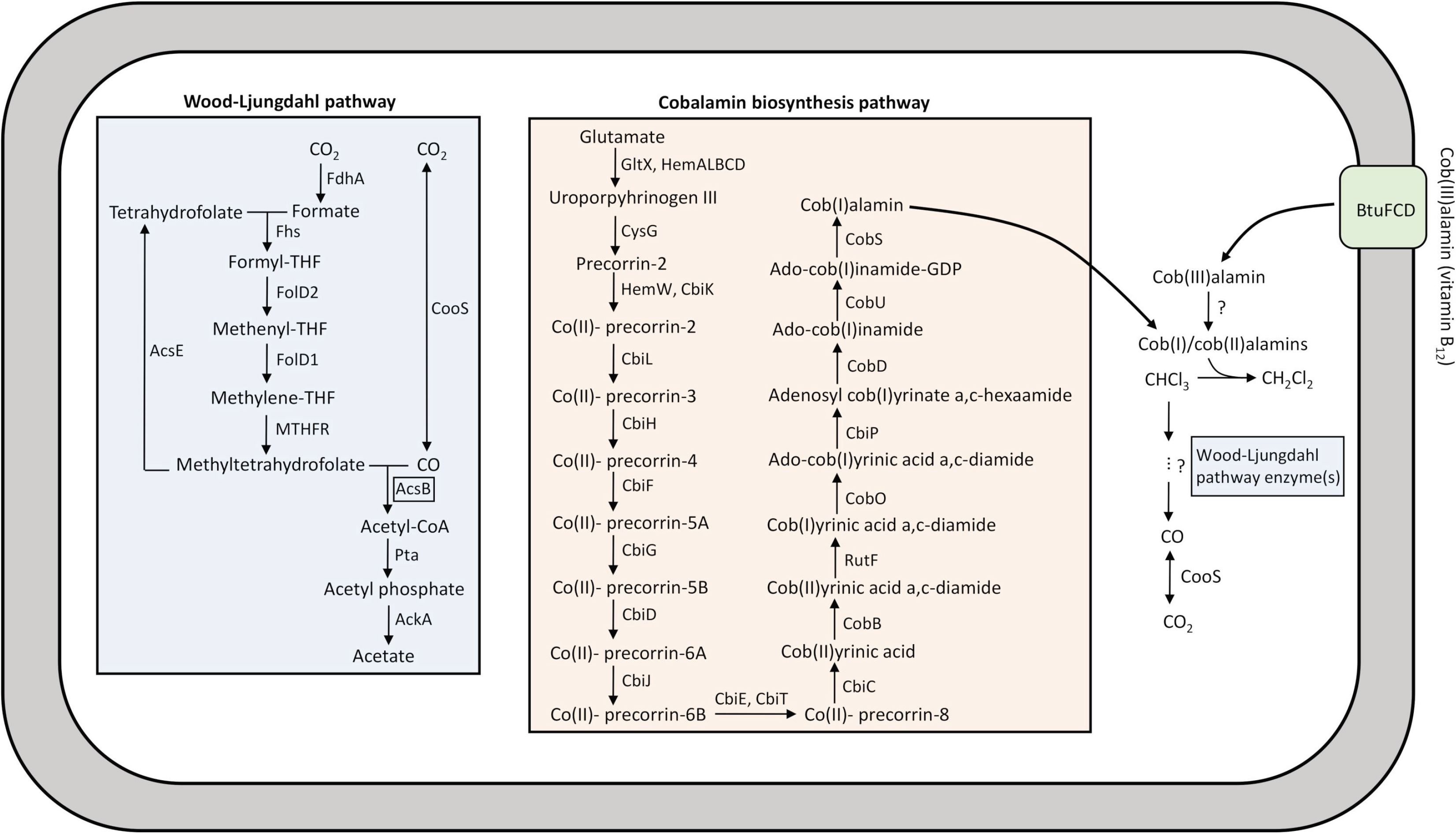
Proposed CF transformation pathway in *Clostridiales* presumably mediated by Wood-Ljungdahl pathway enzymes and cob(I)/cob(II)alamins that are biosynthesized *de novo* or transported from the extracellular environment. The gene encoding AcsB (enclosed in a square) was not found in the metagenomes.

Hypersaline lakes are among the major sources for VOX emissions on Earth (16). In this study, we showed the potential of sediments from pristine hypersaline Lake Strawbridge for CF transformation in cultures with moderate salinity (5%) and alkaline condition (pH 8.5). One possibility is co-metabolic CF transformation through reductive dechlorination and net hydrolysis (Fig. 5) likely mediated by acetogens and/or fermentative bacteria as was proposed in studies using *Acetobacterium woodii* (51), *Clostridium* sp. (not known for its acetogenic potential) (25) and *Pantoea* spp. (23). The MAGs obtained in this study contained most of the genes encoding the *de novo* cobalamin biosynthesis pathway, however, addition of external vitamin B_12_ was essential for enhanced reductive dechlorination and net hydrolysis of CF to DCM and CO. Cobalamin biosynthesis potential has been reported in metagenomic analyses of hypersaline aquatic and terrestrial environments (52, 53). Enhanced CF transformation in the presence of cobalamin indicates an important role of cobalamin not only in fulfilling important ecosystem functions such as carbon processing and gene regulation, synthesis of nucleotides and amino acids (54, 55), maintaining an abundant and diverse microbial community (53), but also potential roles in reducing CF emission to the atmosphere.

## Materials and Methods

### Sampling site

Duplicate sediment cores of approximately 24 cm length and 4 cm internal diameter were collected from Lake Strawbridge (LS, 32.84°S, 119.40°E) and Lake Whurr (LW, 33.04°S, 119.01°E) in Western Australia (Fig. S4). Sediment cores were taken by pushing a polypropylene tube into the sediment. The top and the bottom of the tube were immediately closed with rubber stoppers after pulling the core from the sediment. The sediment samples were transported at 8°C to the Laboratory of Microbiology, Wageningen University & Research, The Netherlands.

### Physical-chemical analysis

The sediment cores were cut into a top (0–12 cm) and a bottom (12–24 cm) layer in an anoxic chamber filled with an atmosphere of N_2_/H_2_ (96 : 4%). Subsamples from each sediment layer were homogenized and subsequently used for physical-chemical analysis and as inocula for enrichment set up. The remaining sediments were kept at −80°C for molecular analysis. Water content was determined by the percentage of weight loss observed after drying the samples overnight at 105°C in an oven followed by cooling down to room temperature in a desiccator. pH was measured immediately and again after two hours using a pH meter (ProLine B210, Oosterhout, The Netherlands) with air-dried sediments suspended in 0.01 M CaCl_2_ solution. Sediment total organic carbon (TOC) was measured using the Kurmies method (56). Low crystalline iron was extracted from 0.5 g wet sediment for one hour in the dark using 25 ml of 0.5 M anoxic HCl (57), and concentrations of dissolved Fe(II) and Fe(III) were quantified using the spectrophotometric Ferrozine assay (58). Major anions including Cl^−^, SO_4_^2−^, NO_3_^−^ and ClO_3_^−^ were analysed using a Thermo Scientific Dionex™ ICS-2100 Ion Chromatography System (Dionex ICS-2100). Major cations including Ca^2+^, K^+^, Mg^2+^ and Na^+^ were measured using inductively coupled plasma-optical emission spectroscopy (ICP-OES, Varian, The Netherlands). Salinity was calculated based on the NaCl concentration (weight/volume) as described before (59).

### Microcosm preparation

Due to dominant presence of halophilic microbes in hypersaline environments (60), and in an attempt to find halophilic microbes capable of CF metabolism, two media were used for halophilic bacteria and archaea enrichment: modified growth medium (MGM) and DBCM2 medium (DBC) (61). The media were boiled and flushed with nitrogen to remove oxygen. Na_2_S·9H_2_O (0.48 g/L) was added as the reducing reagent and resazurin (0.005 g/L) was added as redox indicator. Tris-base (10 mM) and acetic acid (10 mM) were used as the buffer for MGM and DBC media at high and low pH, respectively. The salinity (5–20%) and pH (4.6–8.5) of the media were adjusted to the corresponding values measured in the sediments used as inocula (Table 1, Table S2).

Initial sediment enrichment cultures were prepared in 50 ml serum bottles with 2.5 g wet sediment of either the top or bottom layer of lake sediments and 25 ml of either MGM or DBC medium. The bottles were sealed with Teflon lined butyl rubber stoppers, and the headspace was exchanged with N_2_ gas (140 kPa). CF was added to each bottle at a nominal concentration of 1.25 μmol/bottle. All cultures were set up in duplicate and incubated statically in the dark at 37°C. Of all cultures, the sediment enrichments containing the bottom layer sediment of Lake Strawbridge in MGM with 5% salinity showed better CF dechlorination, and were therefore used for all subsequent experiments. Sediment-free cultures were obtained by sequential transfers of this culture (10% (v/v)) in duplicate in 120 ml bottles containing 50 ml MGM except that peptone was decreased from 5 to 0.5 g/L and yeast extract was decreased from 1 to 0.5 g/L, and glycerol (10 mM) and CF (2.5 μmol/bottle) was added as a carbon source. The sediment-free cultures were used to test the influence of vitamin B_12_ (0.04, 0.4, 0.8, 1.6 and 4 μM) on CF (5 μmol/bottle) dechlorination. Abiotic controls for CF transformation were performed in modified MGM with a decreased amount of peptone (0.5 g/L) and yeast extract (0.5 g/L) and glycerol (10 mM), and amended with 4 μM vitamin B_12_ and 5 μmol/bottle CF, and the same inoculum that was autoclaved at 121°C for 30 min. In a subset of abiotic controls, titanium(III) citrate (Ti(III), 5 mM) or DTT (100 mM) were used as artificial electron donors (62, 63). To test CO_2_ production from CF, ^13^C-labelled CF (99%, Cambridge Isotope Laboratories, Inc., Massachusetts, USA) was used for detecting production of ^13^CO_2_. ^13^CO_2_ formation in the cultures was monitored as outlined below. Cultures without ^13^C-labelled CF were prepared in parallel by supplying 100% non-labelled CF and were used for measuring natural abundance of ^13^CO_2_. The CF dechlorination rate was determined as the disappearance of CF (μmol) per day per liter enrichment culture (μmol/day/L) during the incubation period when dechlorination was stably observed.

Sediment-free cultures for metagenome sequencing were grown in modified MGM with and without addition of 4 μM vitamin B_12_.

### GC analysis

Chloromethanes were quantified from 0.2 ml headspace samples using a gas chromatograph equipped with a flame ionization detector (GC-FID, Shimadzu 2010) and a Stabilwax column (Cat. 10655-126, Restek Corporation, USA). The column was operated isothermally at 35°C. Nitrogen was used as the carrier gas at a flow rate of 1 ml/min. Carbon monoxide (CO), Carbon dioxide (CO_2_) and methane were analysed using a Compact GC 4.0 (Global Analyzer Solutions, Breda, The Netherlands) with a thermal conductivity detector (GC-TCD). CO and methane were measured using a molsieve 5A column operated at 100°C coupled to a Carboxen 1010 precolumn, and CO_2_ was measured using a Rt-Q-BOND column operated at 80°C.

### Isotope analysis

^13^CO_2_ was measured in sediment-free cultures containing 1.25 μmol/bottle ^13^C-labelled CF, 3.75 μmol/bottle non-labelled CF and 4 μM vitamin B_12_, and control cultures contained 5 μmol/bottle non-labelled CF and 4 μM vitamin B_12_. The carbon isotope composition of CO_2_ was determined using gas chromatography combustion isotope ratio mass spectrometry (GC/C-IRMS) consisting of a gas chromatograph (7890A Series, Agilent Technology, USA) coupled via Conflo IV interface (ThermoFinnigan, Germany) to a MAT 253 mass spectrometer (ThermoFinnigan, Germany). Sample separation was done with a CP-PoraBOND Q column (50 m × 0.32 mm ID, 5 um film thickness; Agilent Technology, Netherlands) operated isothermally at 40°C using helium as a carrier gas at a flow rate of 2.0 ml/min. Sample aliquots of 0.1–0.5 ml were injected at split ratios ranging from 1:10 to 1:20. The carbon isotope signatures are reported in δ notation (per mill) relative to the Vienna Pee Dee Belemnite standard.

The amount of ^13^CO_2_ produced from the ^13^C-labelled CF was expressed according to:

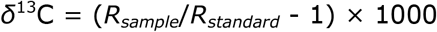

Where *δ*^13^C is the ^13^C isotopic composition (per mil, ‰), *R*_*sample*_ is the ^13^C to ^12^C ratio of the CO_2_ in the sample, *R*_*standard*_ is the international Vienna Pee Dee Belemnite standard (VPDB, ^13^C/^12^C = 0.0112372).

### DNA extraction

The sediment aliquots collected during start-up of the microcosms were thawed and washed three times with 1.5 ml of 10 mM TE buffer (pH 7.0) to avoid interference of the high salinity with the DNA extraction. For each sample, wet sediment (0.5 g) and the washing buffer collected by filtration through a 0.22 μm membrane filter (Millipore, MP, USA) were used for DNA extraction. DNA loss during washing was anticipated, but washing was necessary to be able to extract enough DNA for further analysis (59). DNA was extracted separately from the washed sediment and the biomass collected on the membrane filter using the PowerSoil DNA isolation kit (MO-BIO, USA) following the manufacturer’s instructions. DNA extracts from the sediment and filters were combined for each sample and used for molecular analysis. DNA of the sediment-free enrichment cultures was extracted from 2 ml culture samples using the PowerSoil DNA isolation kit. For metagenome sequencing of the sediment-free cultures, 50 ml of culture was used for DNA extraction using the MasterPure™ Gram Positive DNA Purification Kit (Epicentre, WI, USA).

### Quantitative PCR (qPCR)

The abundance of 16S rRNA genes of total bacteria and archaea, and OHRB including *Desulfitobacterium*, *Dehalobacter*, *Dehalococcoides*, *Sulfurospirillum* and *Geobacter* in sediments (Lake Strawbridge) and the sediment derived enrichment cultures were determined by qPCR. Assays were performed in triplicates on a CFX384 Real-Time system in C1000 Thermal Cycler (Bio-Rad Laboratories, USA) with iQ^TM^ SYBR Green Supermix (Bio-Rad Laboratories, USA) as previously outlined (64). The primers and qPCR programs used in this study are listed in Table S3.

### Bacterial community analysis

16S rRNA gene based bacterial community analysis was performed on sediments of Lake Strawbridge and the sediment derived enrichment cultures. Sediments from Lake Whurr were not proceeded for bacterial community analysis because no CF dechlorination was observed in the enrichment cultures with the sediments of Lake Whurr. The bacterial community analysis was performed as follows: a 2-step PCR was applied to generate barcoded amplicons from the V1–V2 region of the bacterial 16S rRNA genes, and the PCR products were purified and sequenced on an Illumina MiSeq platform (GATC-Biotech, Konstanz, Germany) as described previously (65). Primers for PCR amplification of the 16S rRNA genes are listed in Table S3. Sequence processing was performed using NG-Tax (66). Operational taxonomic units (OTUs) were assigned using uclust (67) in an open reference approach against the SILVA 16S rRNA gene reference database (LTPs128_SSU, version 111) (68). Subsequently, a biological observation matrix (biom) file was generated and sequence data was further analyzed using Quantitative Insights Into Microbial Ecology (QIIME) v1.2 (69).

### Metagenomic analysis

Metagenome sequencing of duplicate sediment-free cultures with and without addition of 4 μM vitamin B_12_ was performed using an Illumina HiSeq platform (PE 150 mode). Fastp v0.19.5 (70) was used for removing adapters and low quality reads. Assembly for binning was done by metaSPAdes v3.11.1 (71) using the option-meta and the trimmed reads. This assembly was used for binning with the Metawrap v1.2 pipeline (docker version) (72). Two bin sets were created from the metagenome samples of duplicate cultures with and without vitamin B_12_ with the bin_refinement module of Metawrap on binners MaxBin2 (73), MetaBat2 (74) and Concoct (75) from the metawrap binning module using the error corrected reads from SPAdes (71). The resulting two bin sets were again run through the bin_refinement module of Metawrap resulting in one bin set containing six bins and unbinned scaffolds. Raw abundance values were taken from the quant_bins module of Metawrap to calculate relative abundances per each culture. A heatmap was created with Python v3.7.3 (http://www.python.org) using pandas and seaborn. Bin quality assessment was performed with CheckM (76). Taxonomic classification of the bins was done by pplacer (77) from CheckM. Annotation of the bins was performed using the Rapid Annotation Subsystem Technology (RAST) (78).

### Sequence deposition

Nucleotide sequences of 16S rRNA genes of bacteria were deposited in the European Nucleotide Archive (ENA) with accession number ERS1165096–ERS1165117 under study PRJEB14107. Raw metagenome sequencing data, primary assembly and assembled bins have been deposited in the ENA under accession number PRJEB32090 (https://www.ebi.ac.uk/ena/data/view/PRJEB32090).

## Acknowledgements

We thank Steffen Kümmel and Florian Tschernikl (UFZ) for their help with the GC/C-IRMS measurement and data analysis, Laura A. Hug and Pascal Weigold for technical assistance during sediment sampling, Mohammad Ali Amoozegar for advice on media selection for halophilic microbes, and the Chemical Biological Soil Laboratory (CBLB) of Wageningen University & Research for assistance with TOC measurement. We acknowledge the China Scholarship Council (CSC) for the support to PP and YL.

---

**Fig. S1** CF transformation by vitamin B_12_ (4 μM) in MGM medium with dithiothreitol (100 mM) (A) or titanium(III) citrate (5 mM) (B) as the electron donor. Points and error bars represent the average and standard deviation of samples taken from duplicate cultures.

**Fig. S2** Quantitative PCR (qPCR) targeting total bacterial and archaeal 16S rRNA genes in the top and bottom layer sediment of Lake Strawbridge and Lake Whurr (A), and sediment enrichment culture and subsequent transfer cultures derived from the bottom layer sediment microcosms of Lake Strawbridge (B). Abbreviation: LS, Lake Strawbridge; LW, Lake Whurr; TOP, top layer (0–12 cm depth); BOT, bottom layer (12–24 cm depth). Error bars represent standard deviations of two (for enrichment samples) or four (for sediment samples) independent DNA extractions, and triplicate qPCR reactions were conducted for each DNA sample (n = 2 (4) × 3).

**Fig. S3** 16S rRNA gene based bacterial community analysis of the sediment of Lake Strawbridge and enrichment cultures. Abbreviation: LS, Lake Strawbridge; TOP, top layer (0–12 cm depth); BOT, bottom layer (12–24 cm depth). Data are shown at phylum level, except *Clostridiales* is shown at order level, and *Halanaerobium*, *Desulfovibrio* and *Bacillus* are shown at genus level. Taxa that were observed at a relative abundance below 1% were summed up and categorized as ‘Others’.

**Fig. S4** Location and overview of Lake Strawbridge and Lake Whurr. The coordinates of the sampling points and the depth profile are shown in the photos. The photos are a courtesy of Christoph Tubbesing from the Department of Geosciences, Universität Heidelberg.

